# Bone mineral density modeling via random field: normality, stationarity, sex and age dependence

**DOI:** 10.1101/2021.02.25.432881

**Authors:** Petr Henyš, Miroslav Vořechovský, Michal Kuchař, Axel Heinemann, Jiří Kopal, Benjamin Ondruschka, Niels Hammer

## Abstract

**Background and Objective:** Capturing the population variability of bone properties is of paramount importance to biomedical engineering. The aim of the present paper is to describe variability and correlations in bone mineral density with a spatial random field inferred from routine computed tomography data.

**Methods:** Random fields were simulated by transforming pairwise uncorrelated Gaussian random variables into correlated variables through the spectral decomposition of an age-detrended correlation matrix. The validity of the random field model was demonstrated in the spatiotemporal analysis of *bone mineral density*. The similarity between the computed tomography samples and those generated via random fields was analyzed with the *energy distance* metric.

**Results:** The random field of bone mineral density was found to be approximately Gaussian/slightly left-skewed/strongly right-skewed at various locations. However, average bone density could be simulated well with the proposed Gaussian random field for which the energy distance, i.e., a measure that quantifies discrepancies between two distribution functions, is convergent with respect to the number of correlation eigenpairs.

**Conclusions:** The proposed random field model allows the enhancement of computational biomechanical models with variability in bone mineral density, which could increase the usability of the model and provides a step forward in *in-silico* medicine.

## Introduction

The structural and intrinsic properties of bone are inhomogeneous, and vary across the multiple spatial and temporal scales and population. It has been documented that bone properties vary at the collagen fibrils level as well as the lamellae level, and naturally vary across anatomical sites [1]. Structural inhomogeneities are related to bone fragility and toughness [2, 3, 4, 5]. *Bone mineral density* (BMD) is widely used to study bone properties. BMD is remarkably inhomogeneous [2, 6], and is connected to bone elasticity and fracture risk [7, 8, 9].

The spatial variation of BMD has previously been analyzed through variograms [10, 11], where the authors attempted to enhance the fracture risk prediction ability related to BMD. Other studies have demonstrated significant correlations between the parameters of BMD variograms and both trabecular bone morphological measures and bone strength [12, 13]. On the other hand, no significant correlation was found between vertebrae strength and variogram parameters [14]. Dong et al. [15] demonstrated that bone elasticity variation at the nano scale can be described as a random field. Due to the remodeling process in bone, stationarity and isotropicity assumptions are likely to be violated, but to the authors’ knowledge, this has never been investigated. Recent studies have emerged describing bone properties as a random field. Desceliers et al. [16] introduced a simplified random field model of cortical bone, but it has not yet been calibrated using clinical data. Another study showed that trabecular structure can be generated by an inverse Monte Carlo simulation on Voronoï cells, which exhibited a good match with trabecular morphology [17]. In the study by Luque et al. [18], a density random field of a trabecular *region of interest* (ROI) was modeled with directionally separable autocorrelation functions based on computer tomography (CT). So far, this study by Luque et al. [18] can be considered the first and also only one that considers density as a random field. However, the conclusions in their study are difficult to generalize to the whole bone because they were derived from a bone sample of small size under stationarity conditions.

Unstable pelvic fractures are difficult to treat and current methods of fixation suffer from a high failure rate [19, 20, 21]. A higher risk of fracture fixation failure is associated with lower mineral density values [22, 23]. In addition, local variance in mineral density has been shown to significantly affect the strength of fixation screws [24]. Regional variance of pelvic bone density is insufficiently described in the literature, although it may play an important role in the study of pelvic fractures. Therefore, the pelvic bone serves as a suitable candidate to demonstrate BMD as a random field in the present study.

### Study Aim & Outline

The present study aims to analyze the spatio-temporal variability in BMD of the pelvic bone and to model BMD as a random field. First, a shape registration algorithm was used to geometrically align CT samples (Shape Registration section). In the next step, the Karhunen-Loève expansion (KLE) was employed to simulate BMD as a random field with Gaussian coefficients, see the Karhunen-Loève Expansion section. The new BMD realizations based on the random field model were validated using average bone mineral density 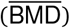, which can be considered a global measure of bone mineral density. Furthermore, what is known as the *energy distance* [25] was computed between the random field of BMD obtained from the CT samples and those generated with the KLE. The energy distance is evaluated *locally* to see how similar the distributions are point-wise, and then also *globally* as an integral measure (Validation Measures section); see the flow chart in Figure 1.

**Figure 1:**
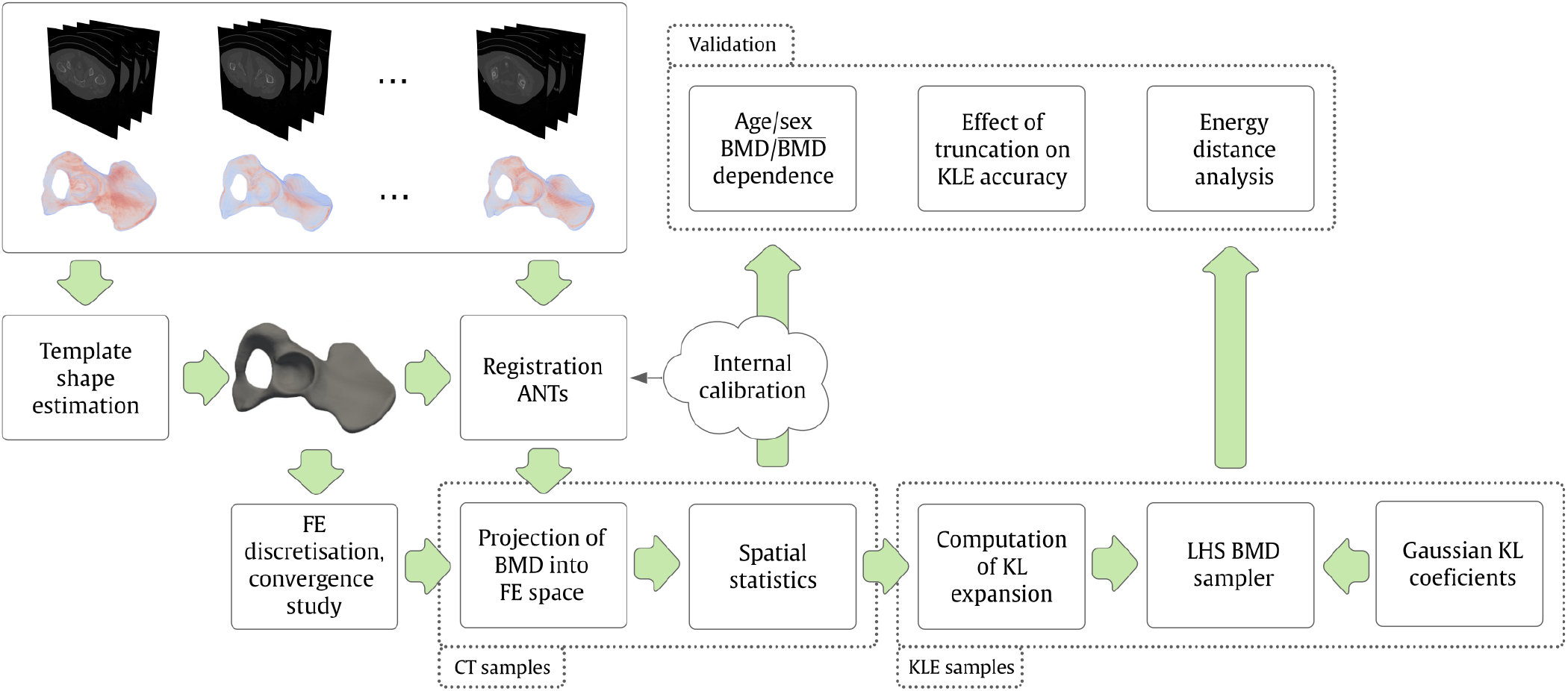
A flowchart of the study.

## Materials and Methods

### CT Data Collection

The anonymized retrospective CT data of 97 females and 88 males were randomly taken from routine examinations performed in the Faculty Hospital in Hradec Králové under ethical approval 202102IO2P. The CT resolution of the dataset was 0.8 × 0.8 × 0.8 mm (Siemens Definition AS+, Siemens Definition 128, both Siemens AG, Erlangen, Germany; 120–130 kV using CareDose, reconstruction kernel 80–90, bone algorithm). The inclusion criteria were as follows: abdominal CT scans, bones without any trauma and, an age range of 20 years or older. Patients who had no record of having undergone a densitometric examination at the time of data collection (2018–2020) were selected. The sample population age per sex is in the range of 22–88 years, divided into 10 bins, where each bin contains more than 5 samples. The pelvic bone geometry implicitly defined by Hounsfield (HU) field was extracted with MITK-GEM interactive segmentation software. First, the rough contours of the bone and background were drawn manually on several slices. Subsequently, the GraphCut algorithm was used to segment the rest of the slices [26].

The CT scans were calibrated internally resulting in BMD [27]. The HU values of air, bone tissue, fat, blood and muscle were considered for internal calibration as shown in Figure 2. Only the right-hand side pelvic bone was considered because no significant difference was identified between the left and right sides.

**Figure 2:**
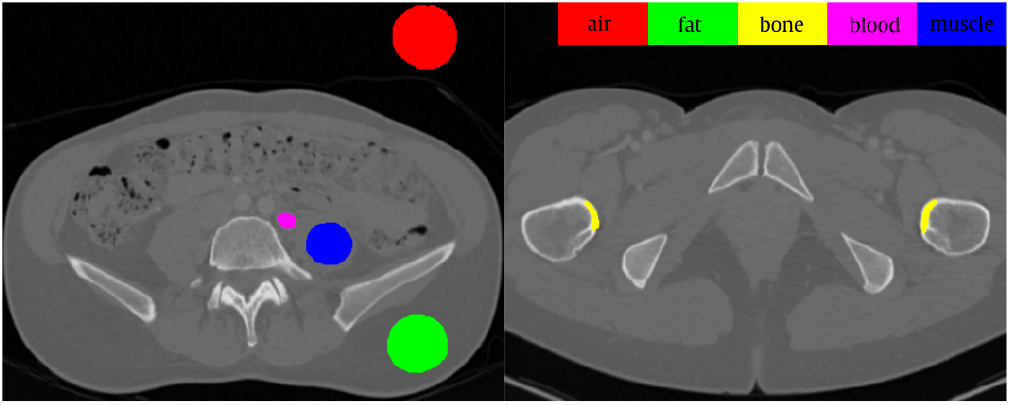
Example of CT slice where HU values of the considered tissues were selected for internal calibration. ROI content (mm^2^): air: 1312; fat: 1109; bone: 160; blood: 92; muscle: 618. Mean HU (standard deviation): air: -1002(7); fat: -90(12); bone: 1233(236); blood: 217(16); muscle: 60(12).

### Shape Registration

The estimation of the random field density requires the universal description of bone locations among all of the experimentally studied bones using a single reference/template bone shape. This is achieved by introducing a fixed metric for spatial or temporal locations per sample to evaluate at. This requirement is violated for bone samples because each sample has a different size and shape. However, bone samples are anatomically and topologically equivalent. This implies the existence of a point correspondence between two shapes under some suitable class of bijective maps and similarity metrics. To find such a correspondence, rigid and affine transforms were realized for the initial global alignment of bones in datasets using the ANTs registration library [28]. Mutual Information (MI) was used as a similarity metric [29]. Then, a non-linear transform was found with the help of the SyN diffeomorphic–based registration algorithm in the ANTs library, see [28, 29]. The similarity of deformed bone shapes was measured with a modified intensity–based criterion called the demons–like metric. This metric provides the best accuracy/speed balance out of all the metrics tested (mean-squared difference, cross-correlation, MI) [29, 28]. In order to minimize registration error, a template bone shape, which is an estimate of the mean sample shape, was estimated according to [29, 30].

### Finite Element Projection of a BMD Field

The template geometry described by an implicit HU field was transformed to a triangulated surface by the marching cube algorithm [31]. The resultant triangular mesh was used to build a tetrahedral volume mesh (fTetWild [32]).

The computer analysis of BMD in the original CT data space is inefficient. Therefore, the BMD is projected into a suitable space with fewer DOFs. In fact, this projection is an approximation of the BMD by piecewise (dis)continuous functions using the least squares method. This approach leads to the minimization of the following functional:

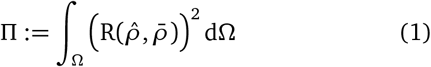

The goal is to find an approximation of the BMD that best represents the original CT data. The residual R represents the difference between the CT BMD value 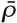 and the approximated value with unknown coefficients 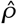:

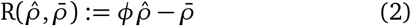

The *ϕ* is FE basis functions evaluated at a given integration point. Substituting (2) into (1) and taking the derivative with respect to coefficients 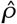, one gets:

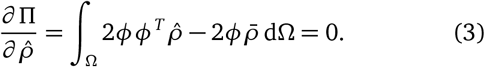

This expression represents a system of linear equations for unknown values of 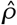:

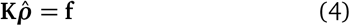

where **K** = ∫_Ω_ *ϕϕ*^*T*^ dΩ and 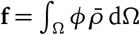. Note that 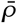 can be noisy, and hence it is evaluated by averaging in a sampling volume of 4 ×4 ×4 voxels in size.

There are two sets of finite element (FE) models used in the present study. The first set consists of validation models. The morphed BMD fields from the dataset were projected onto a discontinuous FE space constructed on the template mesh. All samples in the dataset shared the same geometry domain and finite element space. The correlation matrix of BMD can then be estimated. The FE models in the second set contain BMD fields simulated by KLE on the template geometry. The FE mesh size was estimated based on an auxiliary convergence study where a 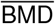 difference between two mesh refinements of below 5% was considered to be converged. The resultant number of degrees of freedom (DOFs) was roughly *M* ≈ 0.7 · 10^6^.

### Karhunen-Loève Expansion

The data set was split into two sets according to sex in order to capture sex differences. Consequently, the relation between age and BMD was analyzed and linear regression was used to separate deterministic trends composing of intercept (sample mean) ***ρ***_0_ and slope ***ρ***_1_ from the data matrix **X**.

The random field ***ρ***(**x**) ∈ Ω is not known explicitly, but only through a set of *N* standardized realizations projected onto the template bone:

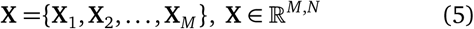

The projected realizations are evaluated at DOF coordinates, from which the matrix of realizations **X** is built. The empirical correlation matrix **C** is estimated as 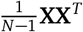. The discretized random field can be viewed as a set of correlated random variables. Sample paths of Gaussian random fields can then be generated by transforming uncorrelated Gaussian random variables into correlated space [33, 34]. One possible linear mapping between the uncorrelated and correlated Gaussian random vectors is via the KL expansion. This expansion involves the eigen-decomposition of the correlation matrix (or the covariance function having the role of a covariance kernel in the continuous version of the KL expansion). In order to compute the KL decomposition of **C**, the associated discrete eigenvalue problem must be solved [35]:

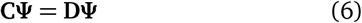

where **Ψ** ∈ ℝ ^*M, M*^ is a matrix of eigenvectors and **D** = diag(*λ*_1_, *λ*_2_, …, *λ*_*M*_) is the diagonal matrix of eigenvalues. The full population of correlation matrix **C** is impossible as it is dense, moreover the rank of the matrix **C** is *N* only and hence we adopt an alternative solution to the above eigenproblem represented by a suitable matrix decomposition. Considering an economical QR decomposition of **X**, the matrix **C** can be expressed:

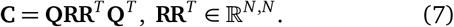

Consequently, the singular value decomposition of product **RR**^*T*^ is computed:

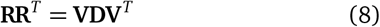

Substitution of Eq. (8) into Eq. (7) leads to:

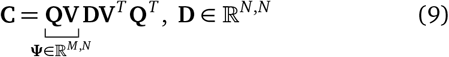

where **Ψ** and **D** are the eigenvector and eigenvalue matrices of **C**. Once the *N* eigenpairs have been computed and sorted in decreasing order *λ*_1_ ≥ *λ*_2_, …, *λ*_*N*−1_ ≥ *λ*_*N*_, the spectral representation of random field ***ρ***(**x**) can be replaced with a truncated discrete KL expansion [35]:

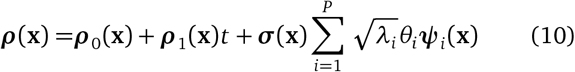

where *θ*_*i*_ is a zero mean, unit variance *i*th Gaussian pairwise uncorrelated variable described by *𝒩*(0, 1), *t* is a time (age) in a range from 22–89 years (from CT data sets) and ***σ***(**x**) is the sample standard deviation.

The truncation in the KLE expressed in Eq. (10) may lead to dramatic computation time savings, since *P* can be considerably less than the order of the correlation matrix (= the number of discretization points), *M*, and also less than the order *N*. An appropriate selection of truncation order *P* can be based on various points of view. The standard way is to control the truncation error in KLE using the decay of the covariance operator’s eigenvalues. The eigenvalues play the role of variances of the underlying uncorrelated random variables *θ*_*i*_, which serve as random coefficients of deterministic eigenfunctions/vectors ***ψ***_*i*_ (**x**). Given this interpretation, one can easily control the total amount of variance represented via the truncated KLE. Since the correlation matrix **C** is positive (semi)definite by definition, the eigenvalues are non-negative and their sum is known in advance. Therefore, the eigenvalues can be sorted from the maximum eigenvalue to the minimum one, along with the corresponding eigenvectors (or eigenfunctions). The gradual sum of the sorted eigen-values serves as an indicator of how much variance is captured by the corresponding subset of eigenmodes. In other words, the expansion can be truncated after taking a subset of *P* dominant eigenvalues (=variables with the largest variance). The number of modes needed to cover a sufficient variability depends on the reach of the autocorrelation function: when the autocorrelation length is high compared to the domain dimensions, usually only a small subset of eigen-pairs is necessary for a given truncation error. Furthermore, it can be shown that the KL expansion is optimal with respect to the global mean-squared error among all series expansions of truncation order *P*. We remark that, in order to achieve convergence, there are restrictions regarding the mesh discretization [33].

The amount of variance captured by the truncated KLE may not be the only criterion for the selection of truncation order, *P*. We also consider stabilization of the energy distance between the generated samples and the required value with *P* as shown in the numerical results below.

In order to generate sample paths of random fields via the KL expansion, a technique for the generation of the underlying standardized pairwise uncorrelated Gaussian random variables *θ*_*i*_ must be employed. Sample paths of random fields generated via orthogonal series expansion directly inherit the quality of sample statistics of the underlying random variables. As shown in [33], utilization of the stratification technique called Latin Hypercube Sampling (LHS) [36, 37] leads to faster convergence of the sample statistics to the target values with increasing number of samples than crude Monte Carlo sampling. Therefore, LHS was used to generate KLE realizations (*n*_sim_ = 300 samples were found sufficient to obtain a converged mean and standard deviation). The LHS generator of pelvic BMD realizations accompanying this paper is freely available on the BoneGen website [38].

### Validation Measures

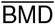 and *energy distance* [25] were considered as validation measures for the proposed BMD random field model. The 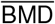 measure is an integral value, defined as

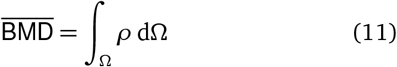

This integral is computed by finite element (FE) discretization. The 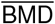 can be considered as the spatial average of the BMD. Since the volume is identical for all samples, it is unnecessary to include a volume denominator in expression (11). Therefore, the 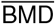 could also be interpreted as a fraction of the bone mass which is formed by mineral content. The energy distance *d* provides a way to measure the similarity between two probability distributions. For two one-dimensional distributions, *u* and *v*, the distance *d* is computed [25]:

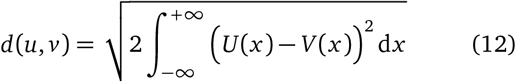

where *U* and *V* are cumulative distribution functions.^1^ Within this study, the expression above describes the spatial distance density over the bone volume, and hence we additionally introduce a global distance measure as well: *D* = ∫_Ω_ *d* dΩ. This spatial integral over bone volume is again computed with the help of FE discretization.

## Results

The mean and standard deviation functions of BMD varied spatially significantly and differed for the cortical and trabecular regions and for both females and males, i.e., BMD random fields were non-stationary in space.

Data analysis for *females* yielded the highest sample mean value of 1.246 (arcuate line, upper third), while the lowest was 0.106 (above the greater sciatic notch). The highest sample standard deviation (std) was 0.191 (top of the acetabular margin) while the lowest was 0.015 (deep to the auricular surface). The BMD normality is considered to be acceptable at the significance level *p* ≥ 0.05, which was fulfilled for 59% of the bone volume. The skewness range is −1.893 (midpart of the anterior margin of the greater sciatic notch) to 7.502 (posterior part of the iliac wing). The negative values corresponding to left-skewed distributions occupy 23% of the volume, while the right-skewed distributions occupy 77% of the volume.

The data analysis for *males* yields the lowest mean value of 0.119 (deep to the auricular surface), while the highest was 1.135 (uppermost part of the arcuate line). The lowest std was 0.016 (in between the iliac wing and the iliac tuberosity), while the highest was 0.218 (top of the acetabular margin). BMD distributions can be considered normal for 54% of the volume, while the rest contained non-normally distributed data. The skewness range is from −1.895 (inferior to the ischial spine) to 6.177 (deep to the auricular surface). The left skewed distributions occupy 17% of the volume, while the rest of the volume was occupied by right skewed distributions. The spatial descriptive statistics are shown in Figure 3.

**Figure 3:**
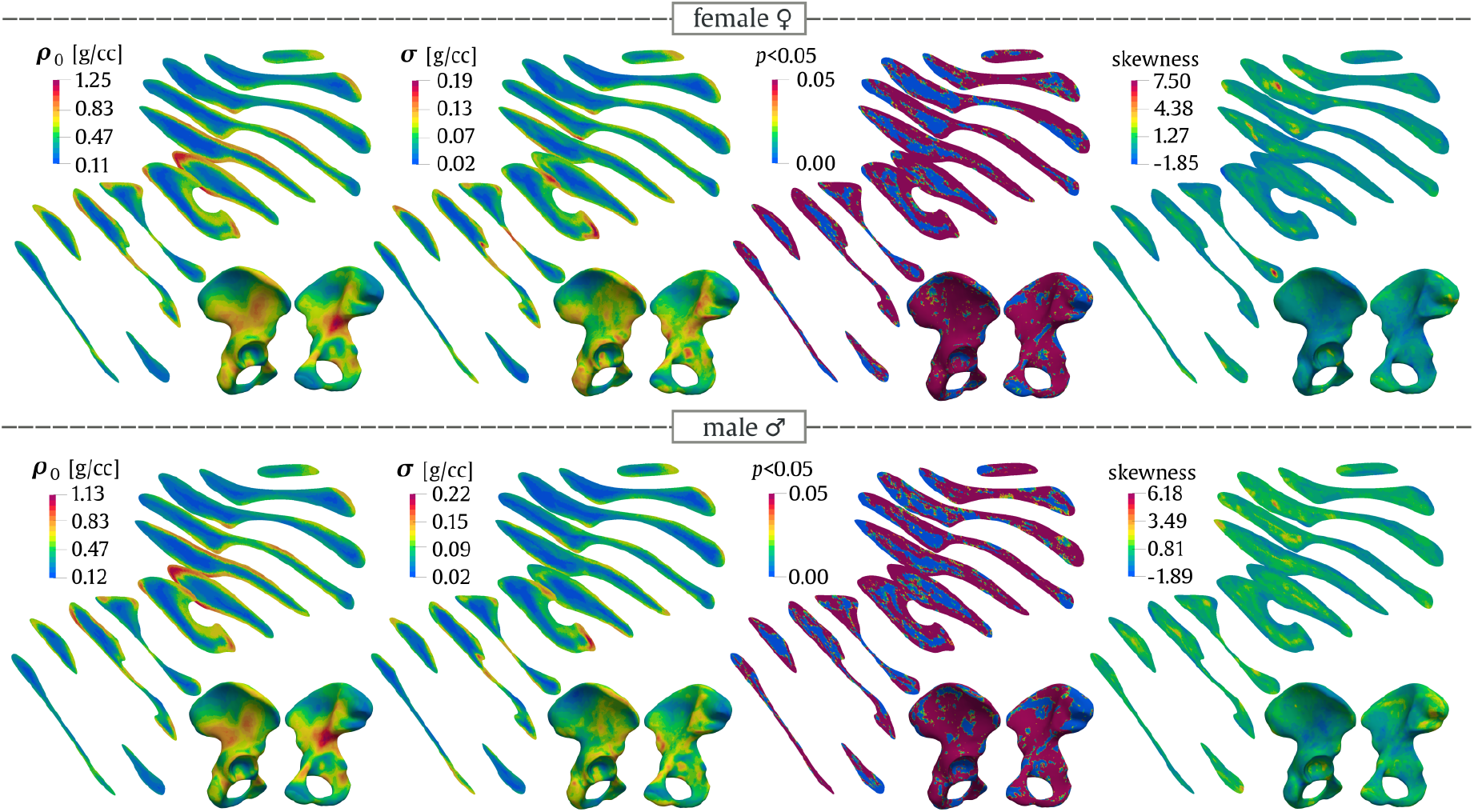
Spatial statistics for BMD comprising three statistical moments for both females and males

### Influence of KLE truncation on the accuracy of random field representation of BMD

The BMD was computed from CT samples and the new samples generated by the KLE with different numbers of eigenpairs. It was found that the most significant eigenvalue explains 32%/36% of the variance in the BMD, and the top five explain 54% of the variance for both females and males. There is no significant statistical difference between the 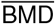 computed from CT- and KLE-based realizations, even with the KLE containing only the most significant eigenpair, see Figure 4.

**Figure 4:**
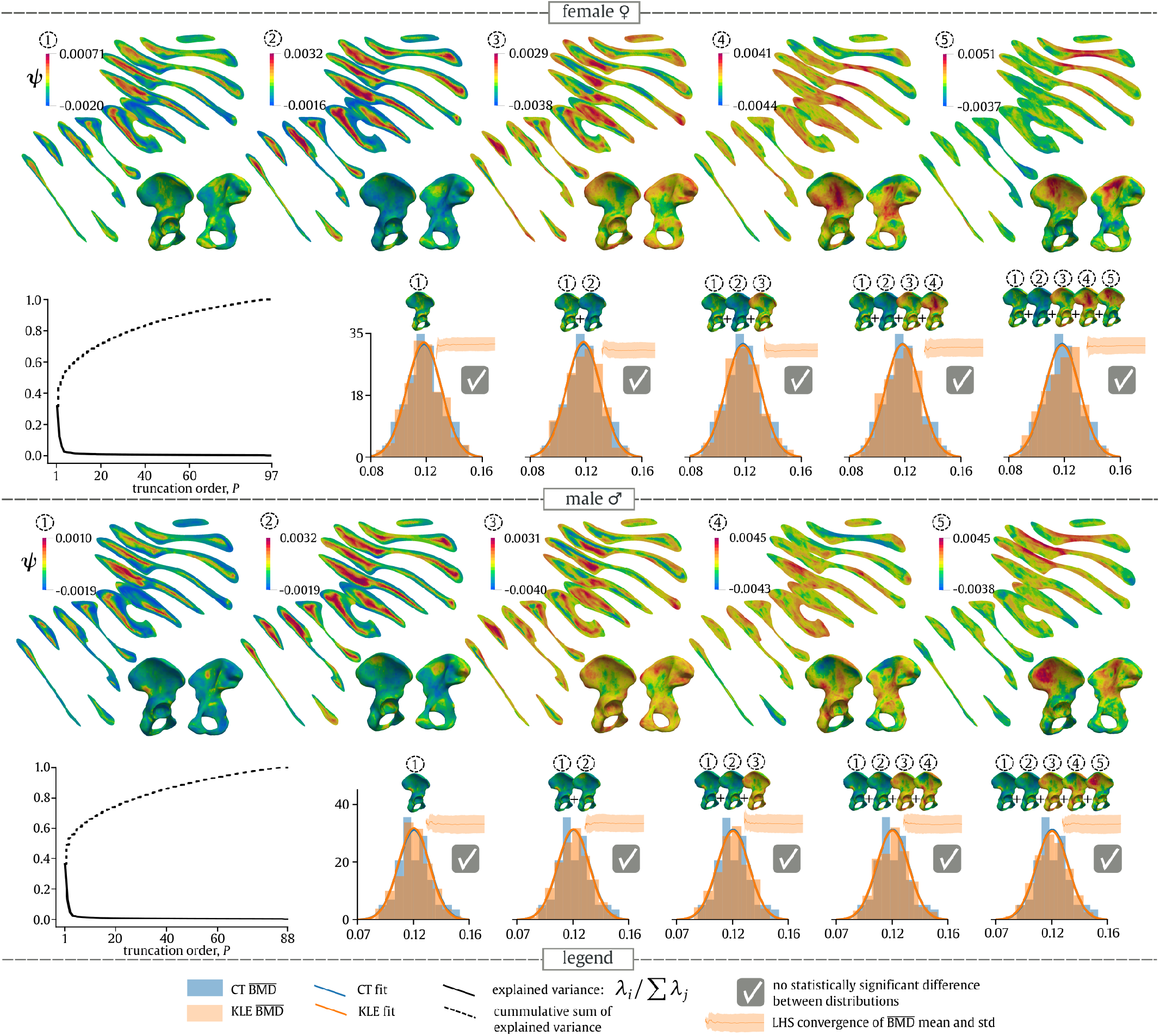
Analysis of explained variance by eigenpairs (*λ*, ***ψ***) and its influence on BMD [g/cc] / 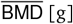 computed by the truncated KLE.

### Age dependence of 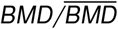

The BMD slope for females varied in range from −5.163 (dorsally to the arcuate line) to 3.269 (above the greater sciatic notch) and from −5.470 (superior-posterior part of the acetabular margin) to 3.625 (anterior third of the iliac crest) [mg/cc/year] for females and males. The BMD is intermediately correlated with age at (*R*^2^ ≤ 0.51) and (*R*^2^ ≤ 0.49) for females and males, respectively. The age correlation was significant at 73% and 56% of volume at a significance level of *p* ≤ 0.05 for females and males respectively, see Figure 5. At 71%/61% of volume, BMD decreased with age for both females and males. The difference in the 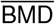 age rate estimated from CT and KLE realizations is 5.57% and 4.71% for females and males, respectively. The difference in standard error was 47% and 55% for females and males. The difference in *R*^2^ is 21% and 50% for females and males; see Table 1.

**Figure 5:**
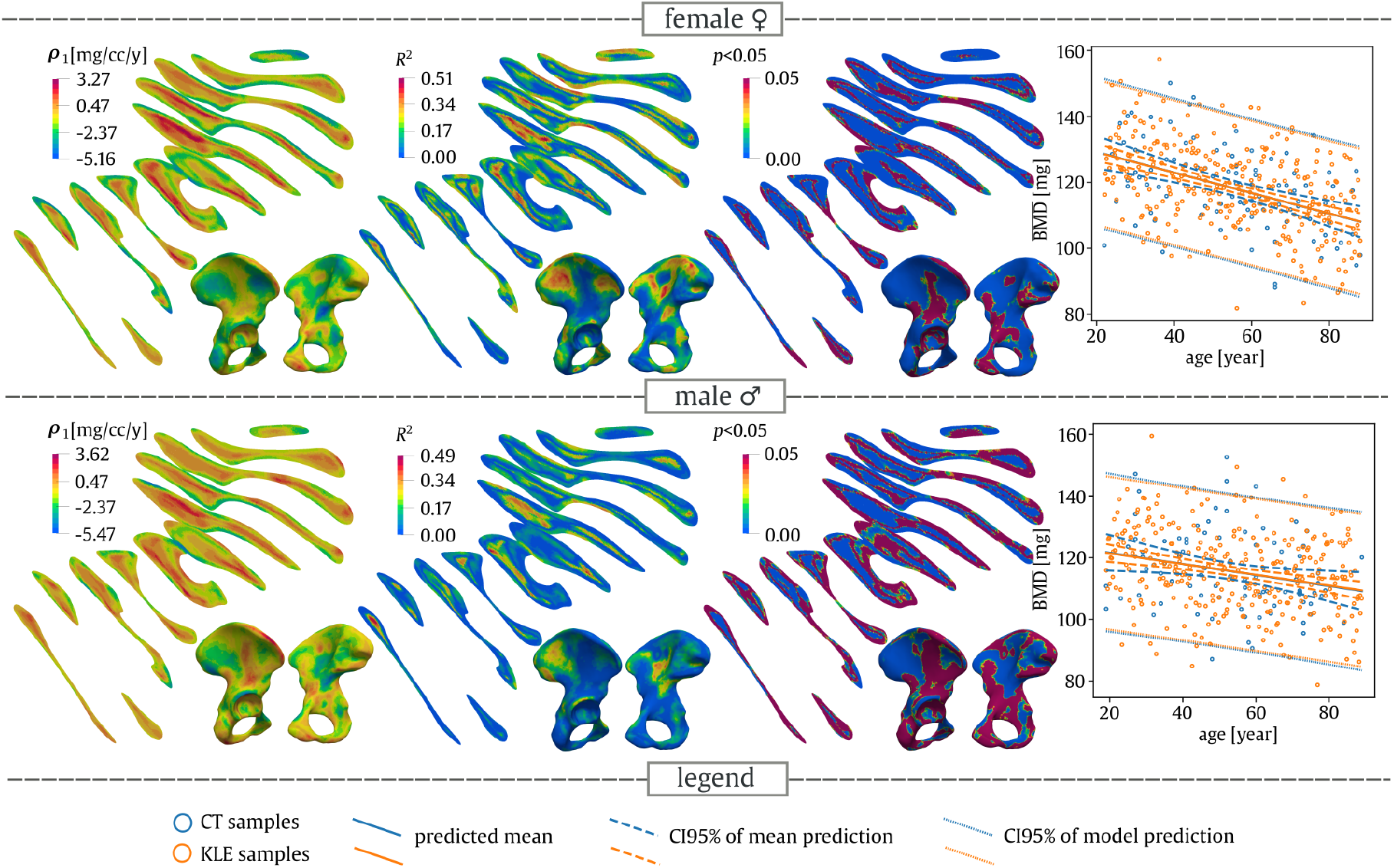
Spatio-temporal evolution of BMD and 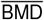.

**Table 1:** Age dependence of 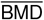 estimated by linear regression for both CT and KLE samples. The KLE samples were generated with five eigenpairs included and LHS design.

| source: | females |  | males |  |
| --- | --- | --- | --- | --- |
|  | CT | KLE | CT | KLE |
| $\overline{\text{BMD}}$ rate [mg/year] | -0.2369 | -0.2501 | -0.1168 | -0.1223 |
| standard error | 0.060 | 0.032 | 0.075 | 0.034 |
| $R^2$ | 0.140 | 0.169 | 0.028 | 0.042 |

### Energy Distance

The minimum/maximum distance *d*_min_*/d*_max_ stabilized after including more than 30 eigenpairs for females. The total distance *D* decreased as the number of included KL pairs increased, and ended up at a value of 7425 for females.

The minimum/maximum distance *d*_min_*/d*_max_ decreased up to the 50^th^ KL pair, and consequently stabilized up to the last KL pair. The total distance decreased as the number of included eigenpairs increased up to a minimum value of 8303. The detailed evolution of energy distance is shown in Figure 6, together with snapshots of selected included eigenpairs. Considering only the first KL pair, there are energy distance peaks at the dorsal portion of the acetabular notch for females and below the anterior inferior iliac spine for males.

**Figure 6:**
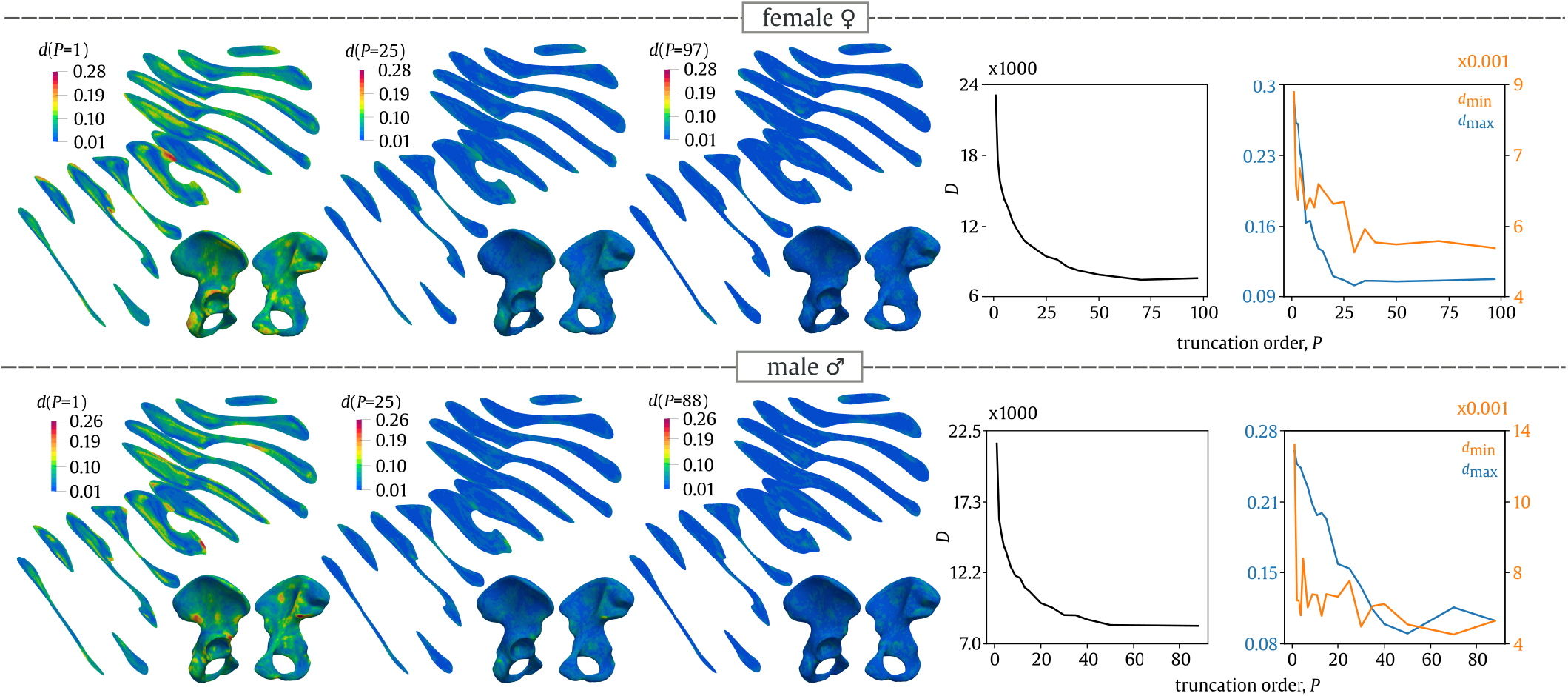
Spatial evaluation of the energy distance composed of spatial functions *d*_min_ and *d*_max_ and the total distance *D* with respect to the number of eigenpairs included. The ratios *d*_min_*/d*_max_ are defined as the minimum/maximum distance over the domain. The minimum/maximum distance location changed with each eigenpair included, which leads to scatter in the convergence plot.

## Discussion

Quantifying the uncertainties in bone mechanical properties originating from a representative population is of paramount importance in order to achieve clinically relevant conclusions and research-informed practice in bone treatment. Due to the complexities of bone shape and the broad individual variations materials, any biomechanical experiments, both real and virtual (for example finite element simulations), should be performed with sufficient sample size. This requirement is often difficult to achieve, and a lack of samples may reduce the potential for research conclusions to be applied to a broad population. We here introduce a random field model for BMD. With this model at hand, one can generate a number of BMD samples respecting population variability and age dependence. The current model allows the replication of the BMD density in a domain, which is a sample mean population bone shape. This step, which aims to separate BMD and shape, allows the analysis of BMD variations at a fixed metric as a random field, but it limits the model’s usability. Nevertheless, shape variations can be considered as a random field as well. The study of BMD and shape variations as random fields, potentially cross-correlated, will form the objective of subsequent studies.

Representing BMD and bone shape using random fields can be considered as a step towards creating a digital twin of bone [39, 40]. However, the next key step is to include osteoporotic changes and analyze their effect on the random field of BMD.

Although patients without a densitometric record were selected and their CT scans were carefully examined by an experienced radiologist, it cannot be excluded that patients with osteoporosis are not present in the considered sample of patients. The patients, although not listed in the database, may have undergone densitometric measurements at another institution or may have been diagnosed with osteoporosis at a later date. In addition, routine CT scans provide limited information about the patient. Furthermore, according to the authors, no information is currently available in the literature on the effect of internal calibration on the accuracy of the T-score used for osteoporosis classification. The above reasons make patient selection by routine CT difficult and must be considered as limitations of this study.

### Spatio-Temporal Dependence of a BMD Random Field

Bone mechanical properties are well known to be age dependent ([41, 42, 43, 44]), and it is likely that the studied random field will also be time dependent. For this present study, only the deterministic part of an age trend was isolated. Generally, a temporal correlation structure can be modeled by the KL expansion but it requires a sufficient sample size per analyzed time period. Knowing the temporal effect on a BMD random field is extremely important and hence it is on the priority list for the authors’ next study.

### Clinical CT Resolution & Calibration

The multi-scale nature of bone could not be considered in detail in the present study. The random field was estimated only at the organ scale based on routine CT data that may not have a sufficient resolution to capture trabecular architecture or the bone cortical shell properly. This issue complicates the estimation of local variations and anisotropy (fabric tensor [45, 46]) of the trabecular network as well as the composite structure of the cortical shell. Although the gradient of the structure tensor might potentially be used to analyze bone anisotropy based on clinical data, this has not been tested in this study [47]. Clinical routine CT is known to distort cortical density and thickness [48, 49], thereby exceeding a 100%-error in the sub millimeter structure of cortical bone. The effect of insufficient CT resolution may be seen at the central part of the iliac wing, where the thickness of the trabecular bone layers is minimized and prone to partial volume effects; this is likely to affect the random field. In some cases, even a fenestration may be present at this location [50]. It is not obvious how the statistical moments and correlation structure are affected, and a careful analysis should be performed with the help of cortical thickness and the density estimation algorithm introduced in [51], dedicated for clinical CT.

The CT data were calibrated internally, without a phantom, using surrounding tissues [27]. Recent studies have shown that internal calibration can be a full alternative to the gold phantom standard [27, 52, 53]. However, various factors that influence internal calibration remain up for debate and therefore caution is in order with regard to achieving accuracy and robustness. Fortunately, the correlation structure of the mineral density is invariant with respect to any linear calibration. However, the mean and variance of the mineral density can be biased by insufficient calibration. In an extreme case, the calibration curve can be considered a source of uncertainty in the mineral density model.

### Spatial Variation of BMD

We assume that spatial fluctuation of BMD reflects the response of bones to external loading, which causes bone to deform in a complex manner (bending + torsion + tension/compression). The load from the trunk is directed through the sacroiliac (SI) joint to the acetabulum and the femoral head while standing, or through the ischial tuberosity while sitting. Simultaneously, more than thirty muscles and several ligaments are attached to the pelvis, loading the bone with their tension in various directions. Increased BMD in area of the greater sciatic notch, the upper part of the arcuate line and the body of ischium seems to correspond well to weight-bearing load. The relatively low standard deviation in this area could indicate that the weight-bearing load can be considered as a common base load in the population. Even though the force generated by related muscles can be significant, just slight density elevations following the margins of large muscles’ attachments (iliacus, gluteus medius) or isolated peaks for muscles with smaller insertion sites such as the rectus femoris were found. However, an interesting similarity was observed between the high standard deviations and the sites of possible apophyseal avulsions. This could indicate an increased individual localized stress induced by inserted muscles or ligament insertions (anterior superior iliac spine – rectus femoris; anterior superior iliac spine – sartorius; ischial tuberosity – hamstrings; iliac crest – abdominal wall muscles; ischial spine – sacrospinous ligament and coccygeus muscle). The increased standard deviation at these sites could reflect variations in physical activity and other unknown effects. Other sites with increased standard deviation, i.e., the superior acetabulum and anterior margin of the auricular surface, are typical of osteophytes.

### Age Evolution of Bone Density

Most publications generally assume a gradual reduction in bone mineral density with increasing age [54, 55, 56]. However, it remains unclear whether this is a uniform process for all skeletal sites or whether there might be some region dependence [57, 58, 59]. Moreover, due to the variable sssurface-volume ratio and related bone turnover, local differences between cortical and cancellous bone should be expected [60, 61, 62]. The age changes in cortical BMD can be described by cortical thinning, higher porosity, pore diameter and osteon density [63, 64, 62, 65, 66]. Cancellous bone is affected by trabecular loss. In males this is mostly in the form of trabecular thinning, while in females trabecular disconnection occurs [67, 68, 69, 70]. There is, however, little known about the spatial and age distribution of BMD in human innominate bone, as the majority of studies focus on long bone, vertebral or hip examinations. Our results showed general age dependent cortical BMD decline and, surprisingly, local mild trabecular BMD elevation. The reason is unclear, but it could be connected to higher trabecular mineralization patterns, which correlate with age, as documented in [71]. We found that female BMD is more sensitive to age. The BMD decreases with age in more than 68%/58% of the volume of bone for females/males. The 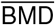 decreases faster for females (51% faster than for males).

### Correlation Structure of BMD

In the present study, a non-parametric approach to the generation of new realizations of BMD has been demonstrated. This approach was based on input CT data, and the next step is to determine parametric correlation kernels, which could represent the correlation structure in time and space. It is unlikely that a simple stationary random field model for whole bone is achievable for several reasons: Bone forms a geometrically highly complex structure, and Euclidean distance is unlikely to be able to properly capture bone topology [72]. Moreover, due to the adaptation processes that bone undergoes, there might be spatially dependent anisotropy in the correlation structure, and the distance metric will be spatially dependent. Finally, multiple latent variables coexist, for example the adaptation process, geometrical influences and other metabolic variables [73]. Together, these variables are very likely to cause long correlation distances, as seen in Figure 7. The identification and separation of these latent variables is difficult due to the limited information available from CT and from patients’ medical records. This will be the topic of a future study. Another question concerns how well the empirical correlation **C** and its eigenpairs represents the true population correlation due to the curse of dimensionality and noise (potentially spurious correlation) [74].

**Figure 7:**
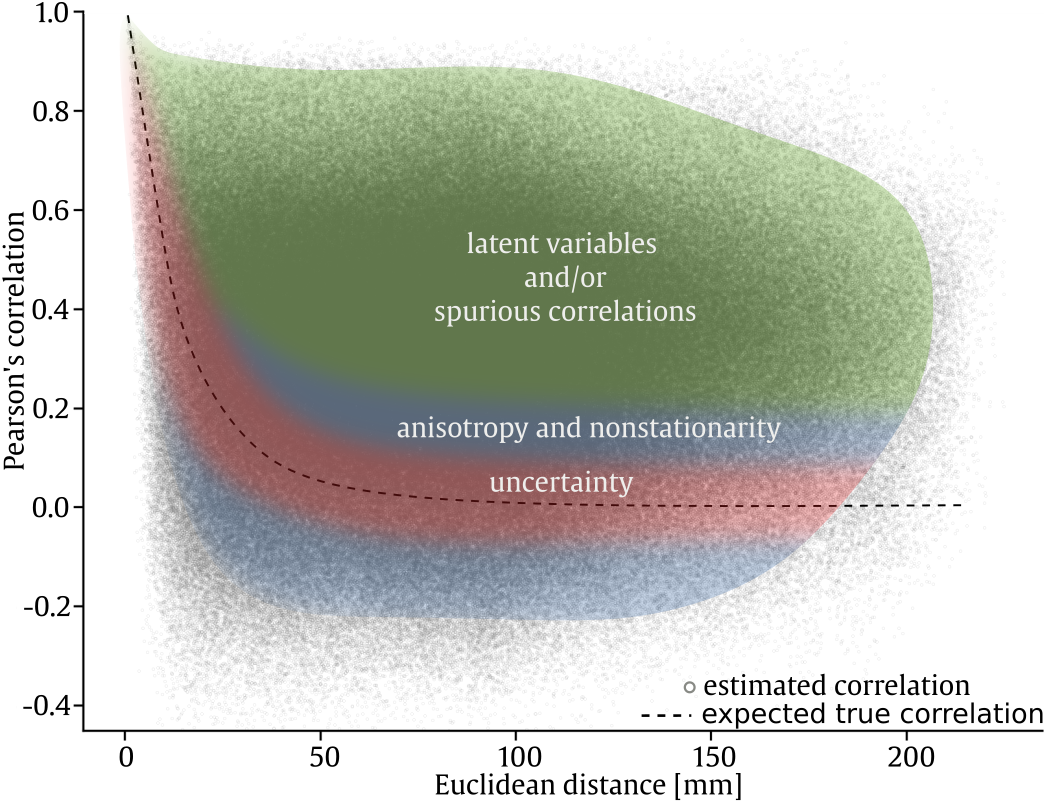
Correlation dependence on distance for a BMD random field for females estimated from CT samples.

### Assumption of Gaussian KL Coefficients

The distribution of BMD is site dependent. There are locations which follow approximately normal distribution, while other locations are slightly left-skewed and significantly right-skewed in distribution as well. The proposed KLE-based model uses uncorrelated Gaussian coefficients, which introduces a certain inaccuracy that is seen in the energy distance metric. The energy metric reveals that the distributions estimated from CT samples and those from the KLE model are different at some locations. It has been shown that five dominant KL coefficients are sufficient for an accurate reproduction of variance in 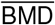. However, the analysis of the energy distance shows that far more KL coefficients (>30) are needed to reproduce the distribution function of the BMD random field. Energy distance is stricter than 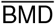 because it directly describes the similarity of BMD distributions. Hence, the energy distance could be a good indicator that local properties such as stress and deformation quantities might not be accurate enough and mean/std estimation might be biased. To improve our model, the identification of (generally non-Gaussian) distributions of KL coefficients should be incorporated into a random field model based on KLE, for example by the iterative algorithm introduced in [75].

### Random Field Model Implementation

The covariance matrix of BMD is dense and large, hence it disallows a common storage representation or the solution of a Fredholm integral equation. Although we partially avoided these difficulties by directly manipulating the data on a discrete level, a more robust approach must be applied, for instance the recent approximation of KL by an isogeometric method [76].

### Comparison with Statistical Shape & Appearance Models (SSM/SSA)

Our method shares the steps of geometry aligment and spectral decomposition of the empirical covariance matrix with SSM/SSA [77, 78, 79], but the meaning and computing of these steps is different. The bone shape aligment is computed on an ROI of whole pelvic bone, allowing the interior to be aligned as well. Our approach uses covariance eigenpairs as bases for generating new BMD realizations. Most importantly, our approach is rather focused on exploring/explaining the spatio-temporal correlation structure, which somehow reflects the (mechano-)biological mechanisms of growth and adaptation [80] in the authors’ opinion.

## Conclusion

The understanding of uncertainties in bone density is of paramount importance to biomechanics in the relation to the understanding of bone mechanobiology, and it should be properly incorporated into computational models. We introduced a random field model describing the fluctuation in bone density via the KLE. The following sub-conclusions can be drawn:

- BMD has a complex correlation structure which cannot be modeled by an isotropic, spatially/temporally stationary Gaussian random field,
- Gaussian KL coefficients allow 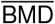 to be simulated accurately,
- the modeled BMD random field allows age dependence of BMD to be incorporated.

## Acknowledgements

The authors acknowledge financial support from project No. LTAUSA19058 provided by the Ministry of Education, Youth and Sports of the Czech Republic. Additionally, the work has been supported by the Czech Science Foundation under project No. 20-01781S.

## Footnotes

1 For empirical distribution functions, the integral is replaced by a sum.

